# Architectural experience influences the processing of others’ body expressions

**DOI:** 10.1101/2023.02.24.529649

**Authors:** Paolo Presti, Gaia Maria Galasso, Davide Ruzzon, Pietro Avanzini, Fausto Caruana, Giacomo Rizzolatti, Giovanni Vecchiato

**Author notes:** **Author Contributions:** P.P., D.R. and G.V. designed the experiment. P.P., G.M.G performed data acquisition and analyses. P.P., P.A., F.C., and G.V. interpreted the results. P.P. drafted the work. P.P., P.A., F.C., G.R. and G.V. revised and wrote the paper. All the authors have contributed to, seen and approved the manuscript.

## Abstract

The interplay between space and cognition is a crucial issue in Neuroscience leading to the development of multiple research fields. However, the relationship between architectural space, the movement of the inhabitants and their interactions has been too often neglected, failing to provide a unifying view of architecture’s capacity to modulate social cognition broadly.

We bridge this gap by requesting participants to judge avatars’ emotional expression (high vs. low arousal) at the end of their promenade inside high- or low-arousing architectures. Stimuli were presented in virtual reality to ensure a dynamic, naturalistic experience. High-density EEG was recorded to assess the neural responses to the avatar’s presentation.

Observing highly aroused avatars increased Late Positive Potentials (LPP), in line with previous evidence. Strikingly, 250 ms before the occurrence of the LPP, P200 amplitude increased due to the experience of low-arousing architectures paralleling increased subjective arousal reports and fixation times on the avatar’s head. Source localization highlighted a contribution of the right dorsal premotor cortex to both P200 and LPP.

In conclusion, the immersive and dynamic architectural experience modulates human social cognition. In addition, the motor system plays a role in processing architecture and body expressions proving how the space and social cognition interplay is rooted in common neural substrates. This study demonstrates that the manipulation of mere architectural space is sufficient to influence human behavior in social interactions.

**Significance Statement:** In the last thirty years the motor system has been recognized as a fundamental neural machinery for spatial and social cognition, making worthwhile the investigation of the interplay between architecture and social behavior. Here, we show that the motor system participates in the others’ body expression processing in two stages: the earliest influenced by the dynamic architectural experience, the latter modulated by the actual physical characteristics. These findings highlight the existence of motor neural substrates common to spatial and social cognition, with the architectural space exerting an early and possibly adapting effect on the later social experience. Since mere architectural forms influence human behavior, a proper spatial design could thus facilitate everyday social interactions.

## Introduction

The interplay between spatial and social environment is a fundamental aspect of daily life (1, 2). The awareness that the amount of time we spend indoors could significantly influence human behavior moved neuroscientists to explore human responses to the built environment, which can be considered the prototypic field for studying the interaction between space and social cognition (3–5). Previous studies demonstrated that the brain contains multiple, plastic, and dynamic space mappings accomplished by fronto-parietal networks characterized by visuomotor properties, mainly described in non-human primate studies and neglect patients (6–10). These cortical regions engaged in space coding partially overlap with networks devoted to action and intention understanding, possibly indicating a functional binding between spatial and social processing (11, 12).

Several studies demonstrated that static architectural features modulate cerebral regions devoted to emotion perception (13–15), and that the motor system is involved in processing affordable architectural transitions (16, 17). From a theoretical point of view, Djebbara et al. provided a psychobiological framework describing the role of the pulvinar in integrating sensory processes, further affecting the higher visual cortex and the related cortico-cortical connections leading to sensorimotor responses integrating environmental features with attention and behavior (18). In addition, Jelic et al. proposed the enactive approach to studying architectural experience, emphasizing the motor system’s role and motivational factors as constituents of the body-architecture interactions (19). Overall, it is recognized that the built environment fundamentally impacts human well-being at multiple temporal and spatial scales, affecting the prevention and containment of infectious diseases (20).

However, despite the increasing number of works in the field (21, 22), the presence and interactions among individuals, and their movement within the architectural space have been neglected so far, failing to provide a unified view of architecture’s capacity to broadly modulate social cognition, such as the perception of other’s body expressions.

In this regard, a large body of evidence has shown that body expressions convey affective information, playing a fundamental role in social interactions (23, 24). For instance, cortical correlates of emotional body expressions perception show increased P200 when observing emotional rather than neutral body postures, pointing to greater attention to socially relevant cues (25). The observation of high-arousing body postures also generates higher Late Positive Potential (LPP) amplitude than low-arousing ones (26, 27). The modulation of such event-related potentials (ERPs) reflects a change in the level of exogenous attention captured by the stimulus (28) and greater attention allocation to motivationally relevant stimuli (29, 30), at an earlier and later stage respectively.

In natural viewing conditions, different stimulus categories carrying affective information, such as people and backgrounds, may all be relevant and processed together, and these information streams may interact. However, only a few studies focused on the effect of the environment in shaping the mechanisms underlying the processing of body expressions, and none of these consider architectural spaces. For instance, behavioral studies showed that the categorization of bodily expressions depends on the emotional characterization of the environment (31, 32). Only one study describes neural evidence showing that the affective information provided by the environment modulates the processing of body stimuli due to the changing activity of cerebral regions endowed with visual functions and others involved in space and body processing (33).

The present study bridges this gap by linking the judgment of emotional body expressions to the dynamic experience of architecture. We exploited virtual reality to ensure a naturalistic experience and requested participants to judge avatars’ emotional expression (high vs. low arousal) at the end of their promenade inside high- or low-arousing architectures. The use of virtual reality is pivotal since it permits subjects to experience the architectural space in a dynamic and immersive way (34, 35), ensuring the same neurophysiological response as in a real scenario (36). Because the processing of emotional body expressions is typically reflected in the modulation of brain components at different latencies, high-density EEG was recorded to investigate the hypothesis that the dynamic experience of architectural spaces modulates neural responses to the avatar’s presentation, thus affecting the early or late stage of attention. Since spatial attention derives from the activation of brain maps transforming spatial information into motor representations (37, 38), we expect to observe a different involvement of motor regions devoted to attention mechanisms and sensorimotor integration depending on spatial and social conditions.

This study demonstrates that the immersive and dynamic architectural experience influences human social cognition. Behavioral and electrophysiological evidence converge toward the modulation of attentional mechanisms at an early stage of body expression processing due to the dynamic architectural experience. In addition, the motor system plays a role in processing architecture and body expressions, proving how space and social cognition interplay is rooted in common neural substrates. These findings reveal for the first time that mere architectural space is sufficient to influence human behavior in social interactions.

## Results

Participants dynamically experienced a virtual architecture before judging an emotional body posture. They perceived the virtual environment through a first-person moving camera, realizing a virtual promenade, and then judged the arousal level of the virtual avatar’s posture (Figure 1A). This procedure allowed us to create a social context to investigate the influence that a dynamic experience of architecture plays on the processing of emotional body postures. Four virtual architectures were selected from a set of 54 models based on their level (low, high) of perceived arousal, in a cold and warm color, as described in a previous study (34) (Figure 1B). Emotional body postures were selected from a set of 45 stimuli based on their level (low, medium, high) of perceived arousal as described in a previous study (39) (Figure 1C).

**Figure 1.**
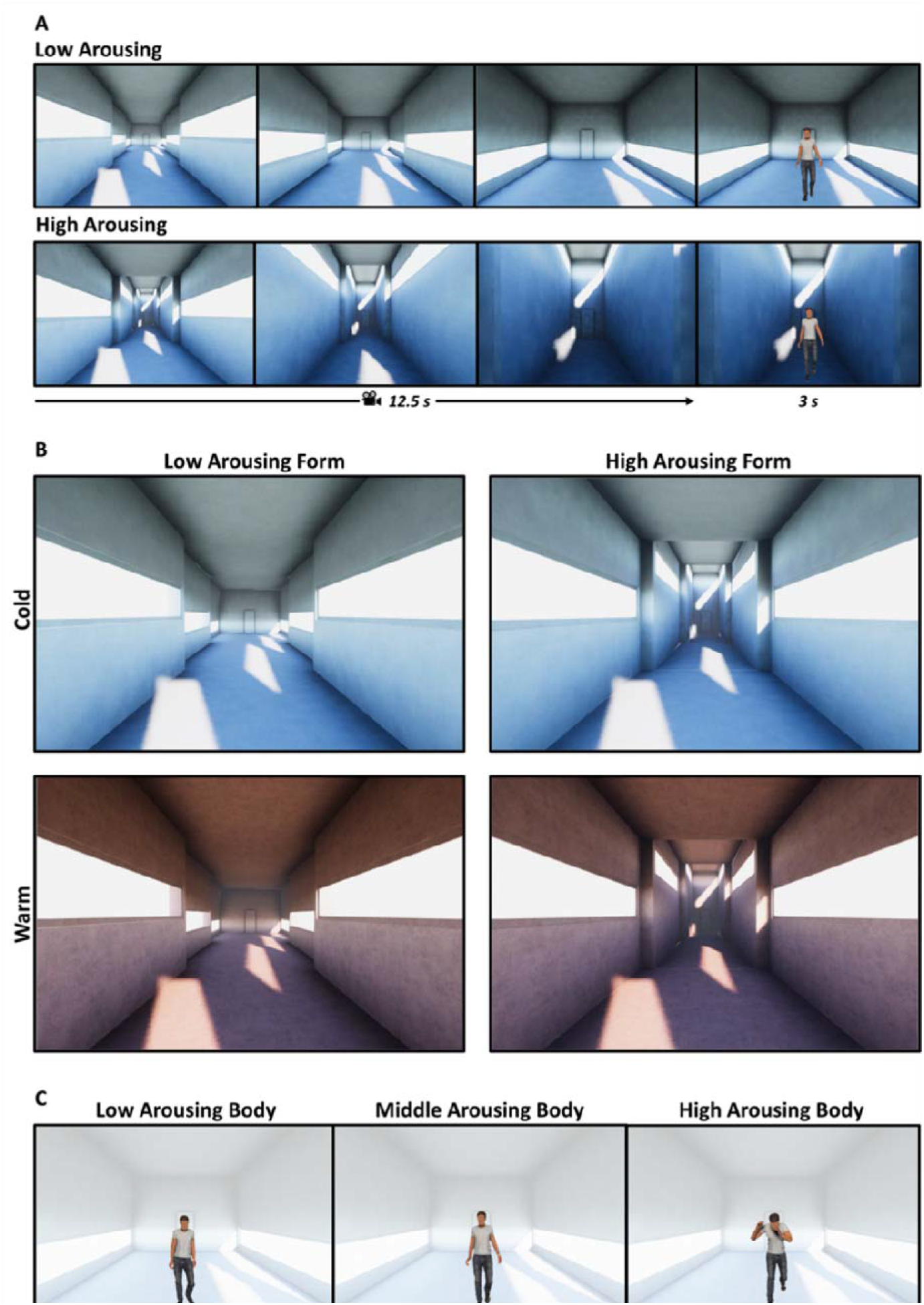
Representation of the experimental trials and virtual stimuli. (A) Schematic representation of two experimental trials. The upper (lower) panels, from left to right, shows three first-person perspectives of the low (high) arousing architecture, corresponding to the participants’ view at the start of the promenade, at the end of the first nucleus, and at the end of the second one. The last frame corresponds to the presentation of the avatar in the third nucleus. (B) Virtual environments with low/high arousing forms (columns) in the cold/warm colored version (rows). (C) Example of avatars with low, middle, and high arousing body posture, respectively. The transparent background is to highlight the body posture and represent the final nucleus of the low arousing architecture.

This EEG study compares ERPs and the corresponding pattern of cortical current density at an early and late stage of the emotional body posture presentation appearing at the end of the virtual promenade. If the processing of dynamical architectural features affects the attention to the avatar at a late stage, we would observe LPP modulations depending on the arousal level of the architecture. Alternatively, if the processing of the dynamical architectural features affects the attention to the avatar at an early stage, we would expect a modulation of the P200 specifically mediated by architectural forms.

### Increased arousal ratings correspond to observation of body postures in low-arousing architectures

The repeated-measures ANOVA returned that participants coherently judged the emotional body postures according to their arousal level (main factor Body: F(2,48) = 115.13, p < 0.001, ηp^2^ = 0.833). Arousal ratings were significantly different among the three levels of avatar’s bodily arousal (low < middle, p < 0.001; low < high, p < 0.001; middle < high, p < 0.001; Bonferroni corrected). These subjective arousal scores were higher within the architectures characterized by low-arousing forms (main factor Form: (F(1,23) = 6.76, p = 0.016, ηp^2^ = 0.227). Instead, the main factor Color (F(1,23) =0.66, p =0.425, ηp^2^ = 0.027) and the interaction Form x Body (F(2,46) = 0.144, p = 0.866, ηp^2^ = 0.006) were not significant. For this reason, the factor Color will not be considered in the following EEG analysis.

### Distinct neural temporal dynamics corresponding to architecture and body characteristics

Figure 2 shows the topographic maps and ERPs related to significant neural activations for the body and form characteristics. We performed a factorial mass univariate analysis in a late and early ERP window. In the late window, the analysis returned a significant modulation of the LPP amplitude related to the arousal level represented by body characteristics. In fact, we report a significant cluster of electrodes for the factor Body (p = 0.006) within the late time interval of 452 - 1000 ms from the avatar presentation. Then, pair-wise comparisons within the main effect Body were conducted through cluster mass permutation tests on the mean ERP amplitudes in this late time window. Specifically, we found a cluster of fronto-central electrodes with higher LPP amplitude elicited by avatars with high-arousing body postures compared to avatars with low-arousing characteristics (p = 0.005). Figure 2A presents the topographic maps of voltage distribution averaged in the late interval for the high- and low-arousing body conditions, showing that the LPP amplitude was mainly located at centro-parietal electrodes. On the right, the grand average ERP of centro-frontal electrodes of the significant cluster is presented, comparing the LPP elicited by the presentation of avatars with high- (blue) vs low-arousing (red) body postures. Also, a different cluster of fronto-central electrodes showed a significantly higher LPP amplitude (p = .014) when avatars had high-arousing body postures rather than middle ones. No significant differences were found comparing avatars with low- and middle-arousing body postures (all p-values > 0.154). No significant clusters were found for the main effect Form (all p-values > .67) and interaction Form x Body (all p-values > .156) in the late window.

**Figure 2.**
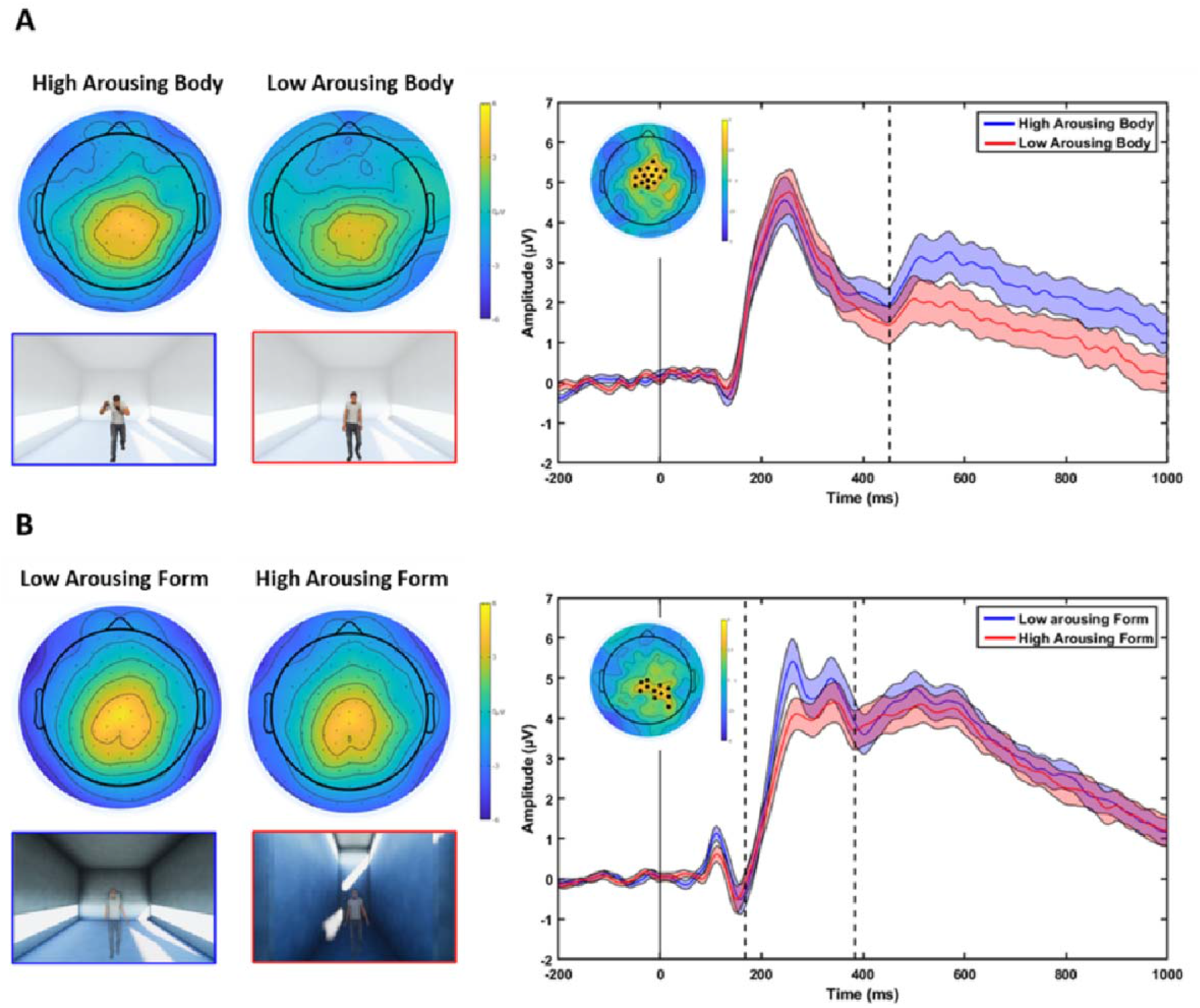
Topographic and ERP activations related to the distinct neural temporal dynamics processing architecture and body characteristics. (A) The left pictures represent the topographic voltage distributions of the LPP (452 – 1000 ms) to the presentation of avatars with high- and low-arousing body postures. The right pictures represent the grand average ERPs for the high- (blue) and low-arousing (red) body posture conditions. (B) The left pictures represent the topographic voltage distribution of the P200 (168 – 384 ms) to the presentation of avatars within low- and high-arousing form. The right pictures represent the grand average ERPs for low- (blue) and high-arousing (red) architecture conditions. Figures within the blue and red frames below the scalp maps highlight the corresponding experimental conditions. The ERPs were averaged across the electrodes defining the significant cluster, highlighted with black dots on the topographic map in the figure inset (colormap codes the t-statistic, cluster-based corrected). The standard error is presented as light shadows of the corresponding color. The significant time interval is defined by back asterisks.

Strikingly, in the early analysis window, the factorial analysis returned a significant modulation of the P200 amplitude during the observation of avatars related to the differences in the architectural forms. In fact, we report one significant cluster for the factor Form (p = 0.023) spanning the early time range between 168 - 384 ms after the presentation of the avatar. Figure 2B shows the topographic maps of voltage distribution and the grand average ERPs of significant electrodes in this early time interval. Specifically, we found a cluster of electrodes in centro-parietal areas with greater activity elicited by the presentation of the avatar within the low-arousing architecture compared to the high-arousing condition. The largest difference between the two conditions was reached around 250 ms after the avatar onset. No significant clusters of electrodes were found for the main effect Body (all p-values > 0.481) as well as for the interaction Form x Body (all p-values > 0.559).

### Common cortical motor activations corresponding to architecture and body characteristics processing

Figure 3A shows the cortical currents density elicited by the presentation of high-arousing body postures and the corresponding statistical cortical map, significantly higher compared to low-arousing ones in the time window between 600 – 660 ms. The dipole with the current density peak within the right dorsal premotor cortex corresponds to a tp = 5.038 (p = 4.24*10^-5^, lower than the FDR corrected alpha threshold 4.29*10^-4^; MNI coordinates: X = 30.6, Y = 7.3, Z = 65). Figure 3B shows the cortical generators of the P200 peak and the significant statistical difference in the right dorsal premotor cortex corresponding to the observation of body postures presented in low-arousing architectures when compared to the high-arousing condition. Specifically, in the 220 – 280 ms time window centered on the P200 peak, we found a significant cluster of activation with tp = 4.861 (p = 6.58*10^-5^, lower than the FDR corrected alpha threshold 2.77*10^-3^; MNI coordinates: X = 18.4, Y = 21.5, Z = 67.1). Overall, findings returned from this EEG study indicate an early-stage modulation of attention mechanisms to the observation of body postures due to the dynamic experience of low-arousing architecture. The activation of premotor areas drives this process.

**Figure 3.**
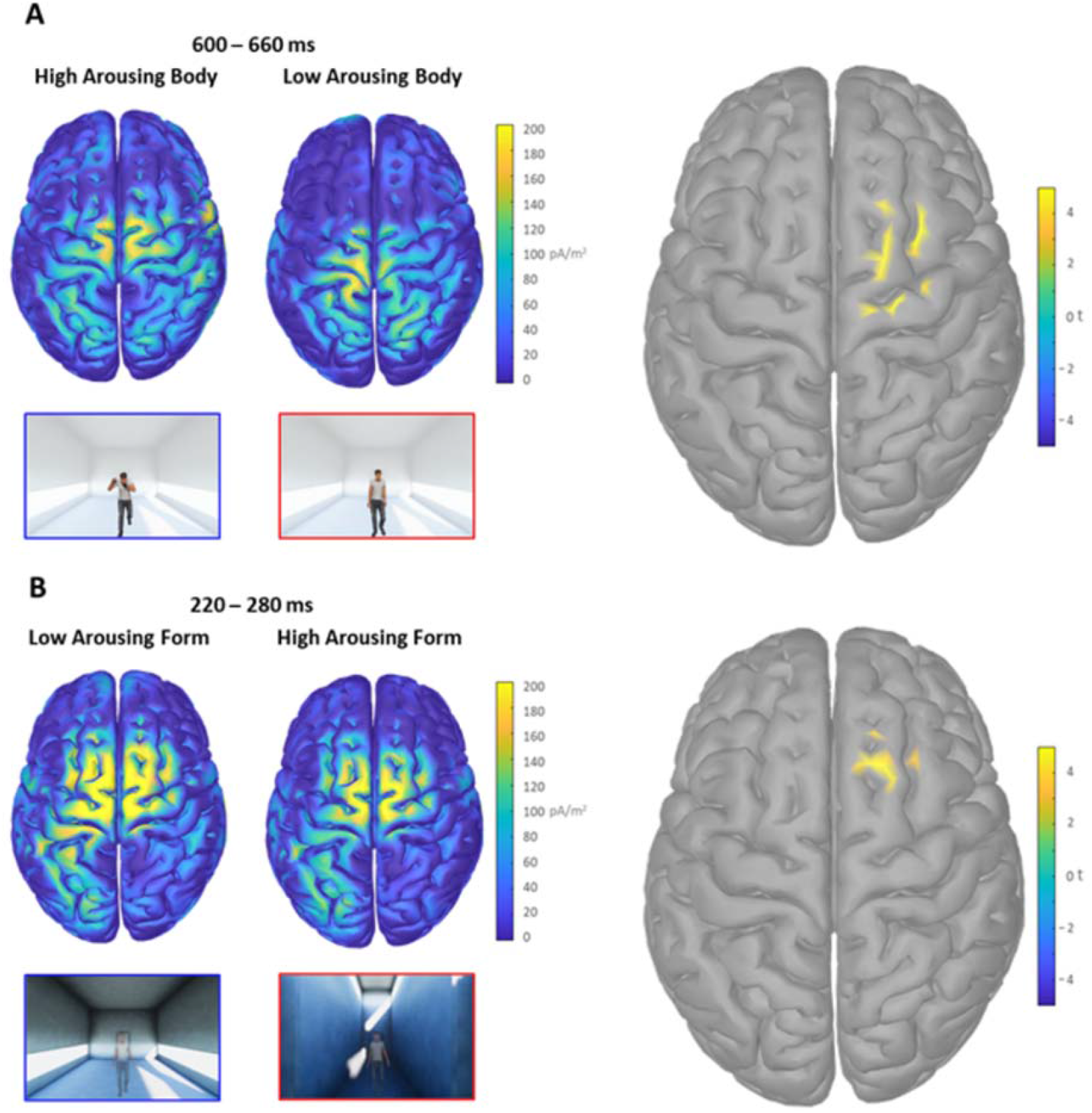
Cortical maps related to the common motor activation for architecture and body characteristics. (A) The left pictures represent the two cortical maps of current density averaged in the 600 – 660 ms interval elicited by the presentation of avatars with high- and low-arousing body postures. The right picture shows the significant dipoles revealed by the corresponding statistical comparison within the cortical map. (B) The left figures represent the two cortical maps of the current density averaged in the 220 – 280 ms interval elicited by the presentation of avatars within the low- and high-arousing architecture. The right picture shows the significant dipoles revealed by the corresponding statistical comparison. The colormaps code the distribution of current density and the corresponding t statistic.

To corroborate these results with covert behavioral correlates of attention, we performed an eye-tracking study to investigate how the fixation times to emotional body postures change according to the different dynamic experiences of architecture. We observed increased fixation times at an early stage of processing on salient avatar’s body districts, such as the head, after the promenade within low-arousing architectures (the eye-tracking study is descried in Supplementary Information). This evidence suggests that the cerebral activations due to architecture characteristics depend on the modulation of attention mechanisms.

## Discussion

The present work explored the interplay between spatial and social cognition by investigating electrophysiological and behavioral reactions to expressive avatars within an immersive and dynamic architectural experience. Reported findings revealed the involvement of late and early attentional mechanisms differently triggered by emotional body expressions and architectural spaces. The observation of arousing body postures elicited increased LPP amplitude, as widely reported in the literature. Strikingly, we found a modulation of the P200 amplitude in response to the avatar presentation, depending on the dynamic experience of different arousing architecture: the more relaxing the architectural experience is, the higher the P200 potential. The source localization highlighted a contribution of the right dorsal premotor cortex to both LPP and P200 generation, pointing to common neural substrates within the motor system processing spatial and body characteristics. Finally, subjective judgments revealed that the avatar’s body was scored as more arousing after the dynamic experience of low-arousing architectures. These findings show for the first time that the dynamic experience of architecture modulates the perception of others’ affective states.

The source analysis revealed that the right dorsal premotor cortex (PMC) is the common neural generator of both LPP and P200, thus involved in the early and late stages of emotional body expression processing. Previous studies suggested the role of the right dorsal PMC in supporting several cognitive functions (40). Specifically, the activity of the PMC reflects the preparation of a motor program to respond to an external stimulus presented in the space, independently from its actual execution. In the present study, the PMC is more activated by the high arousal level expressed by the avatar’s body than the low-arousing one. This result is in line with previous findings showing that high-arousing body postures with socially relevant cues elicit a greater activity of the PMC compared to postures with low arousal levels (25). Strikingly, our results revealed a stronger PMC activity after the dynamic experience of low-arousing architectures at an early stage of the other’s body processing. Hence, we argue that the more relaxing state generated by the architectural experience may foster the preparation of an adaptative social response to the avatar’s body expression, thus eliciting a greater motor readiness in the PMC.

The LPP and P200 are the ERP components sensitive to the late and early processing of the emotional stimulus. On the one hand, the LPP indexes sustained attention on arousing stimuli (29, 41), reflecting the evaluative process of the stimulus significance that may initiate an approaching or aversive response. Our results revealed that the observation of arousing body postures increased LPP amplitude compared with low- and middle-arousing postures, in line with previous research (26, 27). On the other hand, the P200 indexes an early capture of attention, facilitating fast detection of biologically relevant stimuli (28, 42). Strikingly, the dynamic experience of architecture elicited a higher P200 amplitude when observing the avatars within the low-arousing condition compared with the high-arousing one. Previous research found that low-arousing positive states broaden attentional resources (43), while high-arousing negative ones narrow the scope of attention (44). Also, the P200 amplitude is reduced in anxious participants (45, 46) and positively correlates with the availability of attentional resources (47). Hence, we might argue that the modulation of the P200 amplitude reflects a different attentional shift due to the greater availability of attentional resources generated by the relaxing architectural experience.

Redirecting the allocation of attentional resources is fundamental to the attentional control theory, according to which states of anxiety (i.e., a high arousing negative states) impair attentional control by disrupting the balance between the goal-oriented and stimulus-driven attentional system (48, 49). Because the source localization highlighted the involvement of the motor system in the generation of LPP and P200, this result could represent the modulation of attentional mechanisms. In fact, Rizzolatti and colleagues originally proposed the role of the PMC in attention mechanisms, arguing that attention systems are not separated from those for sensorimotor integration (37, 38). In this view, the modulation of PMC would reflect a higher attentional demand requested by specific architectures and body expressions. In parallel, the motor system activity represents a neural signature of embodied cognition, subserving the understanding of spatial and body characteristics (50). Thus, the modulation of PMC in our findings could reflect not the source of attentional demand, rather its target. In other words, higher-order attentional centers could attune the PMC, requesting its higher or lower activity according to the spatial and social experience of the participant.

Finally, subjective arousal ratings revealed that participants perceived the avatars’ body posture as more arousing after the dynamic experience within the low-arousing architecture compared to the high-arousing condition. This result could reflect a conceptual adaptation effect. The virtual promenade within low- or high-arousing architectures represents the adapting stimulus that biases the perception of the arousal expressed by the avatars’ body posture in the opposite direction. Considering the different nature of the two stimuli, the generation of a conceptual adaptation effect is only possible when the adapting and the target stimuli share some perceptual mechanisms (51, 52), reflected here in the common activation in the PMC.

Notably, the difference in the neural activity - depending on both body and architectural conditions - also corresponds to a different pattern of eye movements on the avatar’s body districts as convergence towards the role played by the motor system in integrating attention mechanisms and sensorimotor information (see Supplementary Information).

Over the last few years, researchers have already described the architectural experience in terms of sensorimotor integration, pointing to the modulation of sensorimotor brain areas depending on architectural affordances (16, 17), as well as reflecting the involvement of the motor system during the processing of architectural elements within the surrounding space (14, 53). In this study, the presence of an emotional body expression adds a social component to the architectural experience, which has been neglected so far. Our findings showed that the involvement of motor-related brain areas depends on the spatial experience. Such evidence describes how the architectural space influences the processing of others’ affective states.

## Materials and Methods

### Participants

We recruited 24 participants (26.66 ± 4.02 years, 12 female), satisfying the sample size returned by the power analysis. All participants were naïve to the purpose of the experiment and had a normal or corrected-to-normal vision, with no history of psychiatric and neurological disorders. The local ethical committee approved the study (Comitato Etico AVEN), which was conducted according to the principles expressed in the Declaration of Helsinki. Each participant provided written informed consent before the experiment.

### Stimuli

Virtual architectures were selected from a previously validated database (34), choosing the 2 architectural forms (in a cold and warm colored texture) in which its dynamic experience maximally generated either a low- or a high-arousing state in the participants. Also, an empty environment was designed as a control condition. Avatars’ body postures were selected from a validated database (54), choosing 10 different postures for each level of arousal (low, middle, high).

### Experimental setup and procedure

After reading the written instructions, the HTC Vive Pro Eye head-mounted display (HMD) was comfortably arranged over the participant’s head. Each experimental trial started with 500 ms of static observation of the architectural space from the entrance. Afterward, participants made a straight virtual promenade of 12.5 s crossing the first two nuclei of the architecture (34). Then, participants remained steady for 750 ± 250 ms, and finally a virtual avatar appeared in the middle of the scene for 3 s. Then, participants judged the arousal level expressed by the avatar’s body posture. To this aim, a grey panel was presented reporting the following sentence: “this person looks in a … state” ranging from “Deactivated” to “Activated”. Participants judged the avatar’s arousal by using the Vive Controller. The experiment consisted in 150 trials divided into 6 blocks. The first and the last blocks comprised 15 trials each, where 5 body postures with low-, middle- and high-arousal were presented within the empty environment. Conversely, 30 body postures (10 for each arousal level) were randomly presented within the low- and high-arousing architectures in the central blocks. At the end of each block, participants were allowed to take the HMD off and have some rest.

### Behavioral data collection and analysis

Participants judged the avatar’s arousal by pressing the joypad trackpad button. The cursor of the corresponding panel moved by steps of 0.0083, ensuring a continuous-like movement within the scale ranging between [0, 1], i.e. from deactivated to activated. Then, participants confirmed their choice by clicking the joypad button. Before any statistical data analysis, we discarded trials with possible dips of attention (2.39% ± 3.24). Arousal ratings were z-score normalized considering mean and standard deviation of the scores provided in the empty scene. These normalized scores were analyzed via a 2×2×3 repeated measures (rm) ANOVA, where the within factors were Form (Low-, High-Arousing), Color (Cold, Warm) and Body (Low-, Middle-, High-Arousing).

### EEG data collection and pre-processing

The EEG was continuously recorded at a sampling rate of 500 Hz (vertex reference) using the 128-channels Geodesic EEG System (Electrical Geodesics Inc., Oregon) and the HydroCel Geodesic Sensor Net, which arrays 19 electrode sensors (AgCl-coated electrodes) in a geodesic pattern over the surface of the head at the equivalent 10–20 system locations. Consistent positioning was achieved by aligning the Sensor Net with skull landmarks (nasion, vertex, and pre-auricular points). We used the Net Amps300 high-input impedance amplifier. Low-noise EEG data were obtained, guaranteeing sensor-skin impedances below 50 kΩ except for the reference one, which was kept below 10 kΩ.

EEG data were imported into MATLAB to perform the following analysis with EEGLAB v2021.0 (55). We excluded the outermost belt of electrodes of the sensor net, prone to show residual muscular artifacts, thus discarding 19 peripheral channels located on the cheeks and nape (56). Data were subsampled at 250 Hz, and the PREP pipeline was performed for line noise removal, identification and interpolation of bad channels, and data re-referencing to the common reference (57). To identify ocular, muscular, and remaining channel noise, we computed the Independent Component Analysis (ICA) on the EEG principal components (PCA) that explained 99% of the data variance (55.50 ± 11.21). To this aim, data were firstly band-pass filtered ([2, 100] Hz) (58), segmented in epochs around the avatar presentation ([-1500, 4000] ms), removing the mean value across the epoch (59), and visually inspected to remove corrupted trials (5.46% ± 8.23). Then, we run the runICA algorithm available in EEGLAB v2021.0 (55). Bad ICs (16% ± 7.98) were identified using the ICLabel toolbox (60).

### ERP analysis

The EEG dataset resulting from the Prep pipeline was band-pass filtered ([0.1, 30] Hz) (61) and segmented in epochs around the avatar presentation ([-1500, 1000] ms). Then, ICA weights were applied to the data and pruned the bad components previously identified. A final bad trial rejection (5.43% ± 6.57) was performed by visual inspection. The ERP analysis was performed using the Factorial Mass Univariate ERP Toolbox (FMUT) (62). Firstly, trials were baseline corrected ([-200, 0] ms). Then, two factorial analyses were performed with within factors Form and Body in two different time windows (0 - 400 ms and 300 – 1000 ms) to investigate ERPs at both early and late processing stages. Corrections for multiple comparisons were performed through a cluster-based permutation approach. Specifically, the significance threshold was set to 0.05, the number of permutations to 10000, and the electrode neighbor distance to 4 cm. The FMUT analysis revealed significant spatiotemporal clusters identifying ERP components. Hence, we finally performed pair-wise cluster mass permutation tests on the mean ERP amplitudes resulting from the significant time windows.

### ERP source analysis

We localized ERP sources by solving the inverse problem with the Tikhonov-regularised minimum norm (63). Statistical analysis was conducted at the source level to unveil the cortical generators of the ERPs that emerged at the scalp level. Specifically, for the P200, we averaged the cortical activity within a 60 ms time window centered on the P200 peak and then compared the conditions low- vs high-arousing Form by computing paired t-test (twotailed) between each dipole. Instead, considering that the LPP is a slow tonic component, we averaged the cortical activity elicited within sliding windows of 60ms each, from 420 to 960 ms. For each time window, we then computed paired t-test (two-tailed) between each dipole, comparing the activity elicited by high- vs low-arousing Body. The significance threshold (alpha = 0.05) was adjusted using a false discovery rate (FDR) approach, as implemented in Brainstorm, to correct for multiple comparisons.

Supplementary Information provides details about Power analysis, Stimuli characterization, Virtual environment, Preliminary data clean up, Source localization parameters.

## Acknowledgments

The present study was supported by a research agreement between Lombardini22 and IN-CNR.

