## Supplementary Information for "Architectural experience influences the processing of others’ body expressions"

### Extended methods of the EEG study

**Power analysis.** The sample size for the EEG study was determined using a power analysis computed through the G*Power software (1), considering the 2x3 within-subject design with the "as in SPSS" option and setting the significance level (α) at 0.05, the desired power (1- β) to 0.95, the number of groups to 1, the number of repetition to 6 and the non-sphericity correction ε to 1. The value of the ηp2 was set to 0.17 based on previous research (2), thus identifying the required sample size of 22 participants.

**Stimuli characterization.** Virtual architectures were selected from a database characterized in a previous study where participants rated the dynamic experience of the architecture on the emotional scales of arousal and valence (3). These architectures were conceived as a combination of three consecutive nuclei where the sidewall distance, the ceiling, and the windows sill height could decrease, increase, or remain constant between consecutive nuclei. Specifically, we selected two different architectural forms associated with the lowest and highest level of arousal. The low-arousing architecture is characterized by a constant ceiling, increasing sidewall distance, and decreasing windows sill. Conversely, the high-arousing architecture is characterized by decreasing sidewall distance, increasing windows sill and ceiling height. Also, these virtual architectures were used in two colors, warm and cold. Finally, we designed a control environment exploiting the custom scene of Unity 3D engine Software (2019.1.0f2), characterized by brown-colored ground and a blue sky, adding grey alternate vertical lines in the middle of the scene at the ground level.
Avatars' arousing body postures were selected from a previously tested database via an online experiment where participants scored the visual stimuli using the emotional scales of arousal and valence (4). Three groups of 10 body postures each were selected, representing low-, middle-, and high-arousal, respectively. Different levels of arousal scores across all three groups were guaranteed ($F(2,27) = 102.93$ $p < .001$), as revealed by Bonferroni corrected pairwise comparisons (low < middle, $p < .001$; middle < high, $p < .001$; low < high, $p < .001$).

**Virtual environment.** The HTC Vive Pro Eye head-mounted display (HMD) realized the highly immersive virtual reality environment. This device is equipped with two AMOLED screens, with a resolution of 1440 x 1600 pixels per eye, a refresh rate of 90 Hz, and a field of view (FOV) of 110°. Furthermore, the HTC Vive Pro Eye includes the Tobii eye tracking system, allowing the recording of eye data via the SRanipal (v1.3.0.9) plugin at a sampling rate of 90 Hz with an accuracy of 0.5° - 1.1° (within FOV 20°) https://www.vive.com/uk/product/vive-pro-eye/specs/. Unity was integrated into the HMD via the Steam VR asset to control the experimental procedure and collect data. The experiment ran on a laptop equipped with Windows 10 Home (64-bit), Intel Core i7-9750H, 32 GB RAM, and the NVIDIA GeForce RTX 2070 graphics card.

**Preliminary data clean up.** Before any statistical data analysis, we discarded trials with possible dips of attention. To this aim, we computed the blink rate for each participant and trial. The outliers of the relative distribution were marked and discarded as corrupted trials. Outliers were those trials corresponding to blink rates exceeding the 1.5 interquartile range above the 75° percentile. Using the degree of eye openness recorded by the Tobii eye-tracking system, a blink was identified with a value greater than 0.4 (1 stand for eye closed and 0 for eye opened) as in (5), considering 300 ms of the minimum distance between consecutive blinks (6). This procedure led to the rejection of 2.39% ± 3.24 trials, thus ensuring a blink rate significantly lower after rejecting these trials with possible dips of attention (paired-sample t-tests: $t(23) = 2.493$, $p = .020$). Then, subjective arousal ratings were z-scored.

**Source localization parameters.** The Tikhonov-regularised minimum norm was implemented in Brainstorm (https://neuroimage.usc.edu/brainstorm/Introduction), computing the cortical current density map with dipole orientations that are normal to the cortex (7). As the forward model, we adopted a Boundary Element Method (BEM) volume conduction model of the head from the open-source software OpenMEEG (8, 9) using three realistic layers (scalp, 1082 vertices, 1 conductivity; inner skull, 642, 0.0125; outer skull, 642, 1) based on the head model provided by the FreeSurfer template (ICBM152). We computed the noise covariance from the concatenated pre-stimulus baseline. Then we aligned the electrode locations to the scalp of the head model. As the source model, we adopted a cortical mesh surface with 15002 vertices available in Brainstorm.

#### Eye-tracking study

To corroborate the EEG results with covert behavioral correlates of attention, we performed this eye-tracking study to investigate how the fixation times to emotional body postures change according to the different dynamic experience of architecture. If the cerebral activations due to architecture characteristics depend on a modulation of attention mechanisms, we would observe increased fixation times on salient avatar's body districts after the promenade within low-arousing architectures.

**Participants.** In the eye-tracking experiment 29 participants were recruited (26.96 ± 3.91 years, 14 female). The sample size was determined using a power analysis computed through the G*Power 3 software (1) considering the 2x3 within-subject design with the "as in SPSS" option and setting the significance level ($\alpha$) at 0.05, the desired power (1- $\beta$) to 0.95, the number of groups to 1 and the non-sphericity correction $\varepsilon$ to 1. The value of the $\eta p2$ was set to 0.15 based on previous research showing how participant's gaze on body postures was modulated by their affective state (10), thus identifying the required sample size of 25 participants.

**Eye-gaze data collection and analysis.** Stimuli, experimental procedure, and virtual environment are the same of the EEG study, as described in the previous sections. The HTC Vive Pro Eye includes the Tobii eye tracking system, allowing the recording of eye data such as the eye openness and gaze origin and direction with an accuracy of 0.5° - 1.1° (within FOV 20°) https://www.vive.com/uk/product/vive-pro-eye/specs/. The eye tracking calibration was assessed centering the HMD at the eye level, calibrating the inter-pupillary distance, and finally asking participants to gaze at a central point that moved sequentially to four consecutive peripheral positions.

The preliminary data clean-up procedure led to the rejection of 2.40% ± 2.27 trials, thus ensuring a blink rate significantly lower after rejecting these trials with possible dips of attention (paired-sample t-tests: $t(27) = 4.648$, $p < 0.001$). Then, subjective arousal ratings were z-scored according to the procedure described for the EEG study.

Data from one participant were discarded due to technical issues during the eye tracking recording, thus we analyzed data from 28 participants We computed fixation times (FTs) during the observation of the emotional body postures over 4 different region of interests (ROIs) identifying the head, the trunk, the arms and the legs of the avatar, respectively (11, 12). FTs were z-scored considering the corresponding ROIs in the empty control scene to avoid that possible low-level features, such as size and relative position of different body parts, that could affect the statistics. Also, we discarded fixations that lasted less than 200 ms because any shorter dwell time is typically considered non-fixator activity due to the presence of saccades and potential loss of signal (13–15).

To analyze spatiotemporal dynamics of participant's gaze behavior, we computed the time spent looking at each ROI within time slices of 100 ms each (16). Then, we compared FT differences between the low- and high-arousing body postures as well as between low- and high-arousing architectures. To this purpose, we performed two separate non-parametric analyses based on Montecarlo statistics (5000 iterations, significance threshold 0.05) comparing FT between such experimental conditions across different ROIs and time bins. Considering the strongly correlated spatiotemporal structure of FTs, we adopted a cluster correction method to control for multiple comparisons. Finally a meta-permutation was performed, running the Montecarlo permutation 20 more times and eventually computing the averaged p-value (17).

**Increased arousal ratings correspond to observation of body postures in low-arousing architectures.** Figure S1A shows the results of the rm ANOVA on arousal ratings. As we reported for the EEG study, despite the emotional body postures were coherently judged (main factor Body: $F(2,56) = 92.046$, $p < .001$, $\eta p^2 = .767$), we found increased arousal ratings to body postures observed after the dynamic experience of low-arousing architecture (main factor Form: $F(1,28) = 5.864$, $p = .022$, $\eta p^2 = .173$). Bonferroni corrected pairwise comparisons revealed that arousal ratings were significantly different among the three levels of the avatar's bodily arousal (low < middle, $p < .001$; low < high, $p < .001$; middle < high, $p < .001$). The main factor Color ($F(1,28) = 2.041$, $p = .164$, $\eta p^2 = .068$) and the two-way interaction Form x Body ($F(2,56) = 1.599$, $p = .321$, $\eta p^2 = .039$) did not result in any significant effect.

**Increased fixation times correspond to observation of body postures in low-arousing architectures.** Figure S1B presents the results of the time-varying analysis comparing FTs between low- and high-arousing body postures across different ROIs and time bins. The Montecarlo analysis returned two significant clusters, one for the legs region between 400 and 2800 ms ($p < 0.001$, cluster corrected) and one for the arms from 1900 to 2600 ms ($p = 0.011$, cluster corrected) after the presentation of the avatar. Specifically, participants spent more (less) time looking at the arms (legs) of avatars with high arousing body postures compared to avatars with low arousing ones. Figure S1C shows the results of the Montecarlo analysis comparing the FTs according to the different architectural experience. We found two significant clusters, one for the head region between 700 and 1400 ms ($p = 0.002$, cluster corrected) and one for the trunk from 1000 to 1600 ms ($p = 0.018$, cluster corrected) after the presentation of the avatar, revealing that participants spent more (less) time looking at the head (trunk) after the virtual promenade within low-arousing architectures.

Paralleling the involvement of early and late attentional mechanisms that emerged in the EEG study, the eye-tracking study revealed different patterns of fixation times on the avatar's body district depending both on the arousal level of the body posture and on the dynamic architectural experience. Specifically, results reveal a late allocation of attention on the arms of the avatars with high- compared to low-arousing body postures, showing that the position of the arms is a relevant cue for the recognition of high arousing states (11, 12). Also, we found that participants drew less attention to the legs of avatars with high- compared to low-arousing postures, probably because this body district is not often identified as diagnostic for intensive emotional states (12, 18–20). Strikingly, results revealed that after the virtual promenade within the low-arousing architectures, participants allocated greater attention to the avatar's head and less to the trunk at an early stage of processing. The head position has been found to provide useful information for emotion recognition (12, 18, 19, 21). Also, the higher attention directed at the head of the avatar could reflect the participants' lower anxiety state generated by the promenade within the low-arousing architecture. In fact, people tend to look at other's faces when they feel comfortable during social interactions (10). Thus, as discussed for the brain activities, we argue that such dynamic architectural experience facilitated the initial reallocation of attentional resources on emotional-relevant body cues.

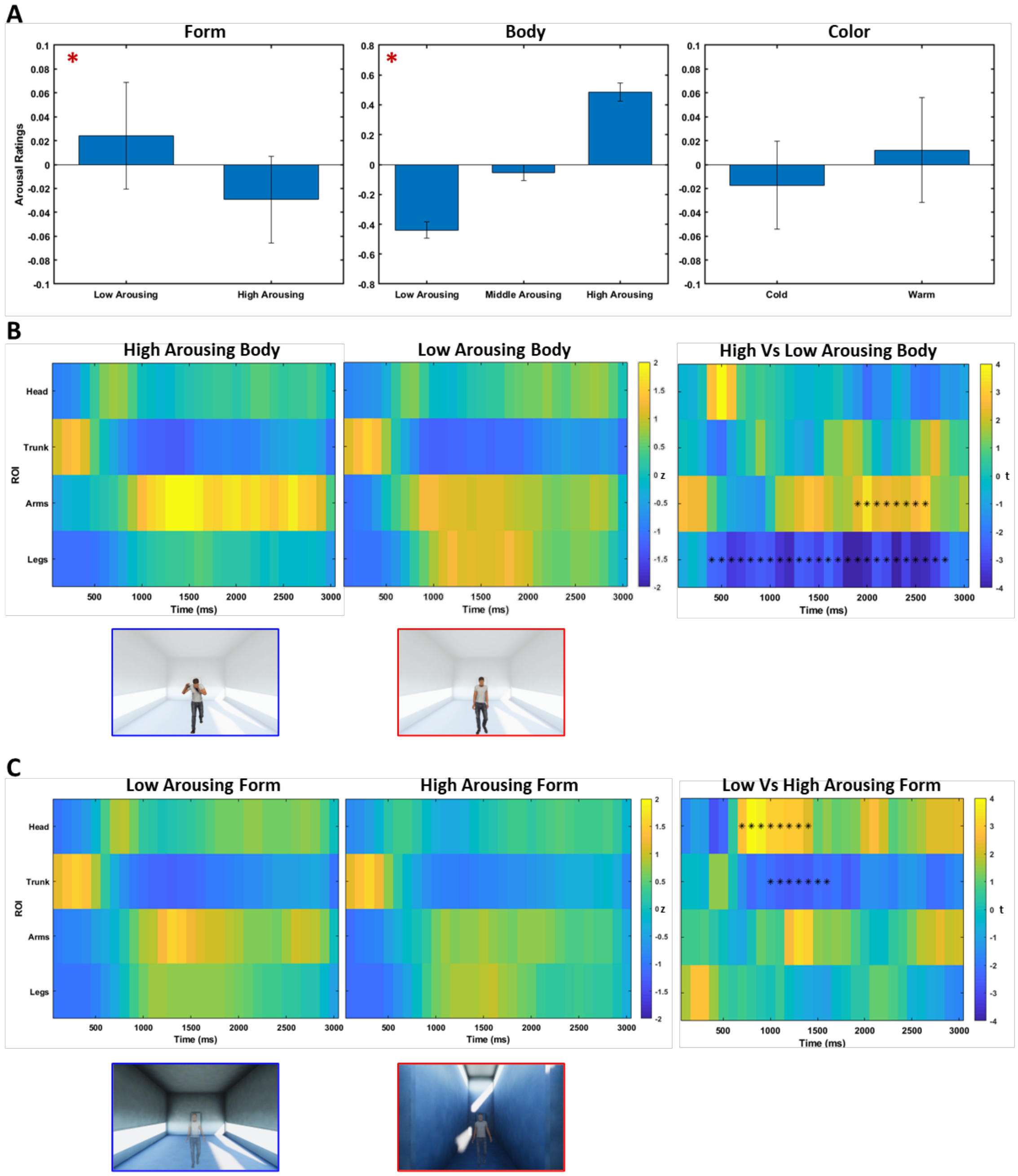


**Fig. S1. Increased arousal ratings and fixation times to body postures after the dynamic experience of low-arousing architecture.** Results of the rm ANOVA on arousal ratings for the main factors Form (on the left) and Body (on the middle) and Color (on the right). Results are presented with their mean and standard error. Significant effects are highlighted by a red asterisk. (B) Time course of fixation times on avatar's ROIs with high- (left panel) and low-arousing (middle panel) body postures. (C) Time course of fixation times on avatar's ROIs within the low- (left panel) and high-arousing (middle panel) architecture. For the left and central panels, the color of each time bin codes the time spent staring at the ROI, z-scored with respect to the empty control condition. In the right panel, the statistical comparison between the two condition is presented: the color of each time bin represents the t-statistic and black asterisks identify the significant cluster related to the head region.
